# Pharmacologically regulated bioorthogonal stabilization domain for regulation of CAR T cells

**DOI:** 10.64898/2026.07.15.738628

**Authors:** Erik Rihtar, Tina Fink, Monika Belak, Rebeka Udvanc, Roman Jerala

## Abstract

Regulation of therapeutic cell response is important for safe and effective therapy, particularly for immunotherapy. Ideally, the regulators should be based on human proteins using compounds that have already been approved for human use. Regulation of protein degradation by small molecules enables fast cellular response and a small genetic footprint of genetic constructs. Here, we present a clinically compatible strategy for reversible pharmacological control of chimeric antigen receptor (CAR) T cell function using human estrogen receptor (ER)–based degron domains and FDA-approved small molecules. By fusing tamoxifen-responsive ER ligand-binding domains to CARs, we generate ligand-inducible ON-switch CARs whose stability, signaling, and effector functions are precisely controlled by 4-hydroxytamoxifen. Furthermore, we demonstrate that ER-tagged CARs can be selectively degraded using the FDA-approved ER-targeting PROTAC ARV-471, establishing a complementary OFF-switch mechanism that suppresses CAR expression and effector function. Notably, these results expand the bioorthogonal ON-OFF switch platform for post-translational control of therapeutic proteins, offering new opportunities to improve the safety and precision of cellular immunotherapies.

## Main

The ability to precisely and reversibly control protein function in living cells remains a central challenge in synthetic biology and therapeutic cellular engineering. In cancer immunotherapy, particularly CAR T cell therapy, excessive or prolonged receptor activity can result in severe toxicities, including cytokine release syndrome and neurotoxicity^1^. Thus, temporal control over CAR activity is critical to balance therapeutic efficacy with patient safety, which still remains a major hurdle to overcome. Consequently, there is a strong interest in developing strategies that enable remote, pharmacological control of engineered T cells. Small-molecule–based control systems are especially attractive because of their ease of administration, predictable pharmacokinetics, and well-characterized biodistribution^2^. One strategy to regulate protein abundance relies on small-molecule–inducible transcriptional control of transgene expression; however, these systems have a slow response, which may be critical for treating adverse events^3,4^. Alternatively, inducible “suicide switch” approaches allow elimination of CAR T cells upon administration of a small molecule, but these strategies permanently ablate the therapeutic cells, which is costly^2^. A more attractive approach is to directly regulate the CAR protein itself, either through conditional assembly of inactive split CAR components^5,6^ or by modulating CAR abundance via ligand-dependent stabilization or degradation^7–9^. Notably, ligand-induced stabilization of CAR receptors has previously been shown to improve CAR T cell phenotype and effector function^10^. The number of available systems suitable for clinically relevant CAR stabilization or destabilization remains limited. Existing destabilized domains include domains like *E. coli-*derived ecDHFR^10^, stabilized by trimethoprim, and FKBP12-based destabilization domains stabilized by Shield-1^9^. These systems suffer from significant drawbacks, including non-human protein origin with associated immunogenicity risks, suboptimal pharmacodynamics, and the lack of FDA approval of the cognate small molecules. These concerns, as well as others, necessitate the development of other bioorthogonal domains stabilized/destabilized with FDA-approved drugs suitable for clinical applications.

An ideal inducible system should employ physiologically inert, titratable, and reversible small-molecule inducers and utilize domains of human origin to minimize immunogenicity. To this end, we focused on the ligand-binding domain (LBD) of human estrogen receptor (ER), which can be modulated by FDA-approved selective estrogen receptor modulators such as 4-hydroxytamoxifen (4-OHT).

Tamoxifen-responsive ER variants have been widely used in biomedical research, most notably the ERt2 domain, a mutant LBD of human ERα harboring three amino acid substitutions (G400V, M543A, and L544A)^11^. These mutations reduce the affinity for endogenous estrogen while preserving sensitivity to tamoxifen, enabling ligand-inducible nuclear translocation of fused proteins. ERt2 has been used in Cre recombinase systems for inducible gene editing in transgenic models^12^. Further engineering of ERt2 through the introduction of additional destabilizing mutations led to the development of the ER50 degron, designed to promote proteasomal degradation in the absence of ligand^13^.

To assess whether ER-based degron tags could regulate CAR surface expression, we engineered Jurkat T cells to express a second-generation anti-CD19 CAR containing 4-1BB and CD3ζ intracellular signaling domains (conventional CAR), or the same CAR fused at the C terminus to either wild-type ERα, ERt2, or ER50 ligand-binding domains (LBDs) (**Figure 1a**). In this design, the addition of 4-hydroxytamoxifen (4-OHT) stabilizes degron-fused CARs, thereby functioning as an ON-switch that enables ligand-dependent CAR T cell activity (**Figure 1b)**. For quantification of CAR stability and expression by flow cytometry, a yellow fluorescent protein (YFP) reporter was incorporated into all CAR constructs. While wild-type ERα did not confer degradation, we unexpectedly found that ERt2 functioned as an efficient degradation domain (**Figure 1c**). Compared to ERt2, ER50 exhibited lower induction and higher basal leakage, likely reflecting its optimization for destabilization of cytoplasmic proteins. Therefore, we focused on the ERt2 domain in subsequent experiments. Importantly, both ERT2- and ER50-tagged CARs were insensitive to endogenous β-estradiol, and CAR stabilization was titratable over a wide range of ligand concentrations, enabling precise control of receptor abundance (**Figure 1e**). We next evaluated the performance of ERt2-regulated CARs in primary human T cells. Treatment with 4-hydroxytamoxifen selectively stabilized CAR-ERt2 in a dose-dependent manner (**Figure 1f**). In co-culture assays with CD19□ Raji target cells, CAR-ERt2 T cells exhibited ligand-dependent cytotoxicity comparable to that of conventional CAR T cells (**Figure 1g**), while conventional CAR activity was unaffected by 4-OHT. Consistent with these findings, antigen stimulation induced robust, ligand-dependent production of IL-2 and IFNγ by CAR-ERt2 T cells (**Figure 1h**). We further examined the reversibility of pharmacological regulation. The inducible expression of CAR ERt2 was detectable approximately 2 hours after ligand addition and reached the maximum after 24 hours, while CAR degradation occurred with a half-life of ∼4.1 hours following 4-OHT washout (**Figure 1i and 1j**). This reversible ON–OFF behavior was also evident at the functional level, as antigen-specific target cell killing could be repeatedly toggled by ligand addition and removal, demonstrating dynamic and reversible control (**Figure 1k)**. Finally, to assess the generalizability of this approach against an alternative target antigen, we generated CAR T cells targeting HER2 antigen. Similarly to anti-CD19 CAR T cells, anti-Her2-CAR-ERt2 T cells exhibited ligand-dependent CAR stabilization (**Figure 1l)**, cytotoxicity (**Figure 1m)**, cytokine secretion (**Figure 1n)**, and T cell proliferation (**Figure 1o)** when co-cultured with HER2□ MCF-7 target cells, with performance comparable to that of conventional CAR T cells. Together, these results establish ER-based degron tagging as a clinically relevant, reversible, and tunable strategy for pharmacological control of CAR T cell activity.

**Figure 1.**
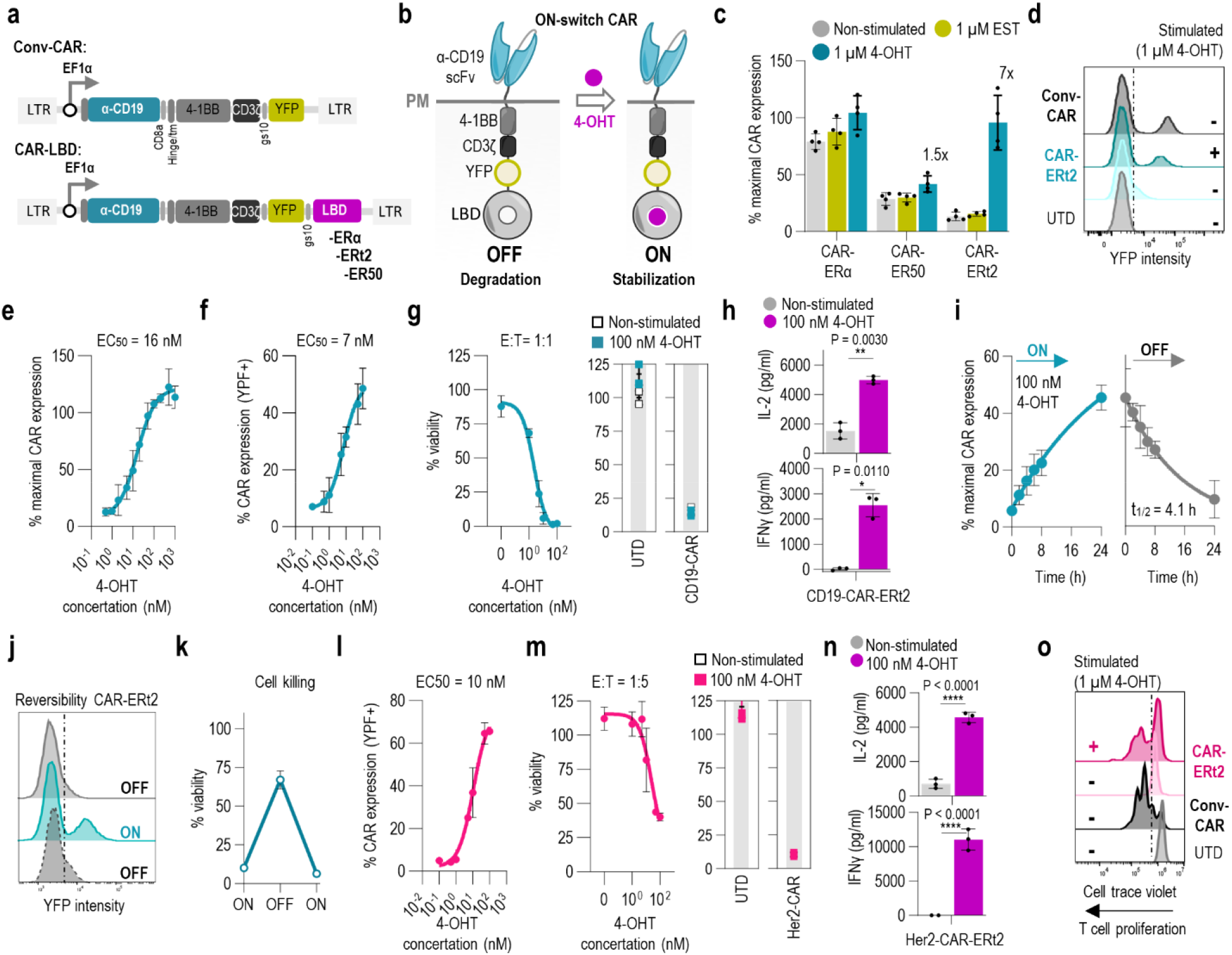
Pharmacological ON-switch CAR T cells. (**a**) Schematic representation of a conventional second-generation anti-CD19 CAR containing 4-1BB and CD3ζ signaling domains, and ligand-binding domain (LBD)–regulated CAR constructs incorporating modified estrogen receptor (ER) LBD variants (ERt2 and ER50). For flow-cytometric detection, YFP was fused to the C-terminus of the CAR. (**b**) Schematic illustrating 4-hydroxytamoxifen (4-OHT) dependent stabilization of LBD-fused CAR constructs. (**c**) Flow-cytometric analysis of anti-CD19 CAR-LBD surface expression in Jurkat cells following 24 h stimulation with estradiol (1 μM) or 4-OHT (1 μM). (**d**) CAR surface expression in Jurkat T cells after overnight stimulation with 4-OHT or vehicle control. UTD, untransduced. (**e**,**f**) Dose-dependent surface expression of anti-CD19 CAR-ERt2 following overnight incubation with increasing concentrations of 4-OHT in Jurkat cells (e) or primary human T cells (f). (**g**) Cytotoxic activity of anti-CD19-ERt2 CAR T cells treated with increasing concentrations of 4-OHT against Raji-CD19 leukemia cells (1:1 E: ratio, normalized to tumor-only control). (**h**) IL-2 (top) and IFN-γ (bottom) secretion by anti-CD19-ERt2 CAR T cells treated with indicated concentrations of 4-OHT after coculture with Raji-CD19 leukemia cells. (**i**,**j**) Flow-cytometric analysis of ON/OFF kinetics of anti-CD19 CAR-ERt2 surface expression at indicated time points following addition or withdrawal of 4-OHT (100 nM). (**k**) Reversibility of anti-CD19-CAR-ERt2 function. Raji-CD19 target cells were treated with 4-OHT (100 nM) for 24 h, washed, cultured for an additional 24 h without 4-OHT, and subsequently re-exposed to 4-OHT for 24 h prior to coculture with CAR T cells. Cytotoxicity was assessed 24 h after addition of CAR T cells. (**l**) Dose-dependent surface expression of anti-Her2-CAR-LBD following overnight incubation with increasing concentrations of 4-OHT in primary human T cells. (**m**) Cytotoxic activity of anti-Her2-ERt2 CAR T cells treated with increasing concentrations of 4-OHT against MCF7-Her2 target cells (1:5 E:T ratio, normalized to tumor-only control). (**n**) IL-2 (top) and IFN-γ (bottom) secretion by anti-HER2 ERt2 CAR T cells treated with indicated concentrations of 4-OHT after coculture with MCF7-HER2 target cells. (**o**) Proliferation of anti-HER2 ERt2 CAR T cells after 5 days of coculture with MCF7-HER2 target cells. Leftward shifts in histogram peaks indicate T-cell proliferation. Data are shown as mean ± SD of triplicate wells. Representative donor shown from two independent donors. All schematics were created using Inkscape (ver. 1.2.1).

We next hypothesized that the ERα LBD could be readily repurposed to enable a pharmacological OFF-switch by exploiting proteolysis-targeting chimeras (PROTACs) that bind ERα and recruit the ubiquitin– proteasome system **(Figure 2a)**. ARV-471 is an FDA-approved ER-targeting PROTAC currently FDA approved for the treatment of ER-positive breast cancer. To test PROTAC-mediated depletion of CAR receptors, we engineered Jurkat T cells expressing a second-generation anti-HER2 CAR fused to the Erα domain. Treatment with ARV-471 induced dose-dependent degradation of the CAR receptor, supporting the feasibility of PROTAC-mediated post-translational regulation of CAR abundance **(Figure 2b and 2c)**.

**Figure 2.**
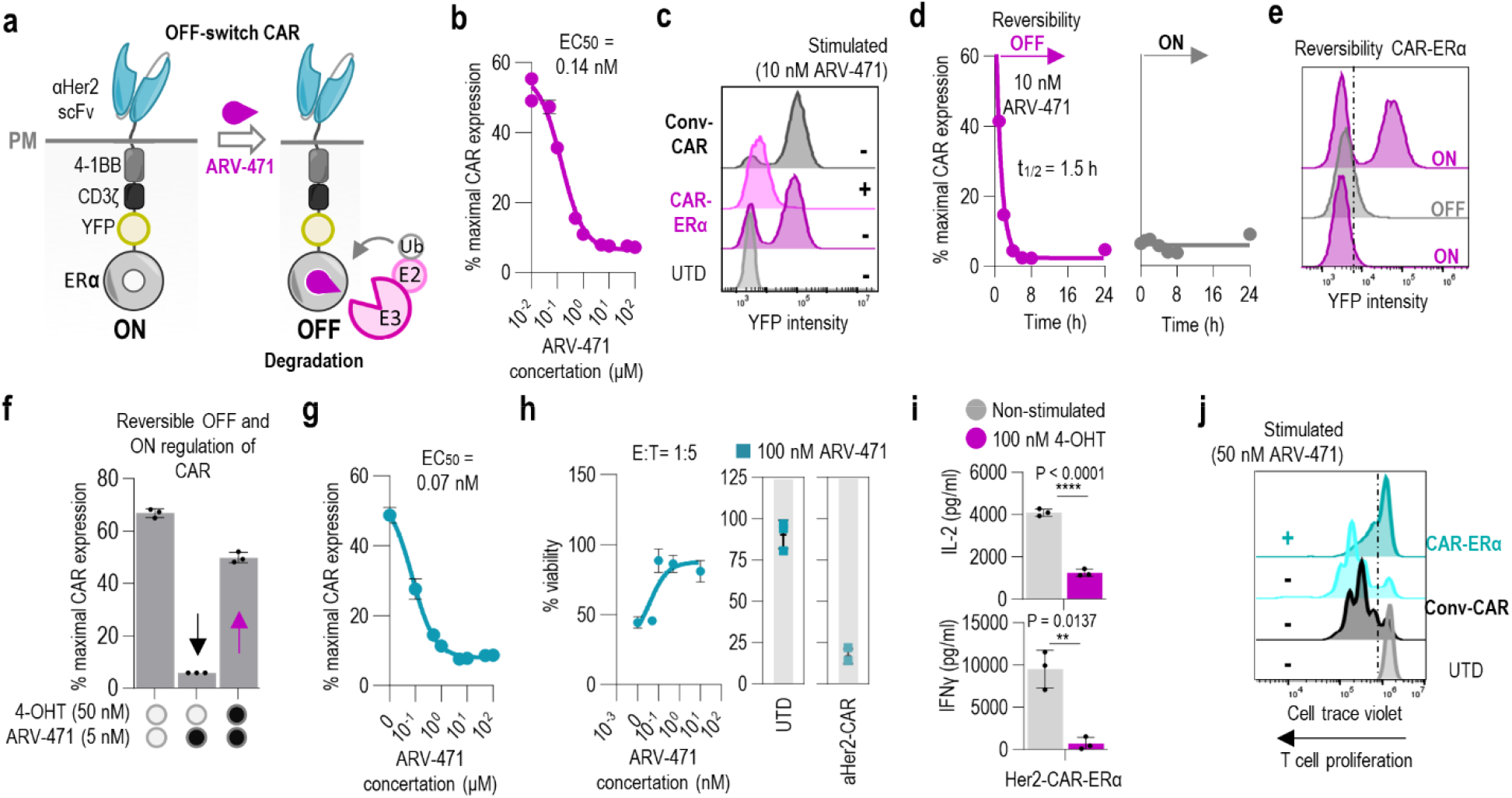
Pharmacological OFF-switch CAR T cells. (**a**) Schematic of the degradable OFF-switch anti-Her2-CAR-ERα, in which receptor degradation is induced by the PROTAC ARV-471. ARV-471 recruits an E3 ubiquitin ligase to the ERα ligand-binding domain, resulting in ubiquitination and depletion of the CAR. (**b**,**c**) Flow-cytometric analysis of surface anti-Her2-CAR-ERα expression in Jurkat T cells following overnight treatment with increasing concentrations of ARV-471. (**d**,**e**) Flow-cytometric analysis of ON/OFF kinetics of anti-Her2-CAR-ERα surface expression in Jurkat T cells at indicated time points following addition (d) or withdrawal (e) of ARV-471 (10 nM). (**f**) Dynamic control of anti-CD19 CAR-ERα using competitive ligands. Simultaneous treatment with 4-OHT and ARV-471 prevents PROTAC-mediated degradation of the CAR receptor. (**g**) Flow-cytometric analysis of surface anti-Her2-CAR-ERα expression in primary human T cells following overnight treatment with increasing concentrations of ARV-471. (**h**) Cytotoxic activity of anti-Her2-ERα-CAR T cells treated with increasing concentrations of ARV-471 against MCF7-Her2 tumor cells (1:5 E:T ratio, normalized to tumor-only control). (**i**) IL-2 (top) and IFN-γ (bottom) secretion by anti-Her2-ERα-CAR T cells treated with indicated concentrations of ARV-471 following coculture with MCF7-Her2 tumor cells. (**j**) Proliferation of anti-HER2 ERt2 CAR T cells after 5 days of coculture with MCF7-HER2 target cells. Leftward shifts in histogram peaks indicate T-cell proliferation. Data are shown as mean ± SD of triplicate wells. Representative donor shown from two independent donors. All schematics were created using Inkscape (v1.2.1).

Kinetic analysis revealed that half-maximal CAR depletion occurred approximately 1.5 hours after ARV-471 addition **(Figure 2d and 2e)**. However, after washout of ARV-471, we did not observe the recovery of CAR abundance. We hypothesized that competitive binding of 4-OHT could recover PROTAC-mediated degradation. Simultaneous addition of 4-OHT and ARV-471 indeed effectively blocked CAR depletion, enabling the reversibility mediated by 4-OHT stabilization **(Figure 2f)**.

We next evaluated functional control in primary human T cells expressing anti-HER2 CAR–ERα. ARV-471 induced dose-dependent depletion of CAR surface expression in primary T cells **(Figure 2g)**. In co-culture assays with MCF-7 target cells, pre-treatment with ARV-471 resulted in dose-dependent inhibition of target cell killing **(Figure 2h)**. Ligand-induced CAR degradation was further reflected in reduced IL-2 and IFNγ secretion **(Figure 2i)**, as well as impaired antigen-driven T cell proliferation **(Figure 2j)**. Together, these results demonstrate that ERα degron tagging enables PROTAC-mediated OFF-switch control of CAR T cell effector functions.

In summary, we describe clinically compatible small-molecule–inducible ON-and OFF-switch domains that enable precise pharmacological control of CAR T cell activity using FDA-approved drugs. Although numerous small-molecule–responsive systems have been developed, the majority are not readily suitable for translational or clinical applications. We demonstrated that nanomolar concentrations of 4-OHT, which trigger weak systemic effects, are sufficient to potently activate CAR T cells. Alternatively, scaffold-based release of 4-OHT or caged 4-OHT^14^ would be possible to selectively activate CAR T cells within the tumor, in case of targeting an antigen that may be weakly expressed also on healthy tissues.

Future studies will be required to evaluate the effects of 4-hydroxytamoxifen– and PROTAC-mediated CAR control *in vivo*. Notably, both 4-hydroxytamoxifen and ARV-471 are clinically approved for the treatment of breast cancer, raising the possibility that these agents could be combined with CAR T cell therapy to achieve synergistic and tunable tumor targeting.

## Acknowledgements

This research was supported by grants from the Slovenian Research Agency (J3-60061, P4-0176, J7-4640, J7-4493, R.J.) and project CTGCT which received Teaming for excellence funding under the European Union’s Horizon research and innovation program Grant agreement ID: 101059842 (R.J.).

## Author contributions

E.R. and T.F. conceptualized the research, E.R. and T.F., designed the experiments. E.R., T.F., R.U., and M.B. performed the experiments on cell culture. E.R. and T.F. analyzed the data and interpreted the results. E.R. drafted the initial manuscript. E.R., T.F. and R.J. critically reviewed, edited and provided feedback on the manuscript. R.J. supervised the work and provided funding.

## Data Availability Statement

All data supporting the findings of this study are available within the article. Any other relevant data or reagents are available from the corresponding author upon reasonable request. Source Data are provided with this paper.

## Methods

### Plasmid construction

All plasmids were constructed using the Gibson assembly method^15^ or traditional restriction/ligation method, and were constructeid in pLVX lentiviral vectors. Second generation CAR was crustucted using CD8 leader signal sequence, Myc tag, anti-CD19 scFv (FMC63) or anti-Her2 scFv (4D5), CD8 hinge, transmembrane and 4-1BB and CD3z stimulatory domains, along with yellow fluorescent protein (YFP) (marker for surface CAR abundance) and ER derived domains (ERa, ERt2 and ER05). Plasmids were aplified in NEBstable competent cells.

### Mammalian cell culture and cell stimulation

The cell line Lenti-X™293T cells (Takara-Bio, LentiX293) was cultured in DMEM medium (Thermo Fisher Scientific) supplemented with 10 % v/v FBS (Thermo Fisher Scientific). At 30–90 % confluence, LentiX293 cells were transfected with a mixture of DNA and PEI (6 µl/500 ng DNA, stock concentration 0.324 mg/ml, pH 7.5). The Raji lymphoma cells (ATCC), Jurkat E6.1 (ATCC, Jurkat) and MCF-7 breast cancer cells (provided by Toni Petan, Jozef Stefan Institute, IJS) were cultured in RPMI-1640 medium (Thermo Fisher Scientific) supplemented with 10% v/v FBS. Raji-fLuc and MCF-7-fLuc reporter cell lines were created by lentiviral transduction and puromycin selection (3 µg/ml) (Thermo Fisher Scientific). All cell lines were cultured in a humidified incubator at 37 °C with 5 % CO_2_. Stock solutions of 4-hidroxytamoxifen (10 mM in DMSO, Sigma Aldrich, Catalog No. H6278), 17β-estradiol (10 mM in 100 % DMSO, Sigma Aldrich, Catalog No. 2834) and ARV-471 (1 mM in DMSO, MedChemExpress, Catalog No. HY-138642), were stored at -20 °C until use. Cells were stimulated by adding the appropriate concentration of the ligand by 1000x dilution of the stock concentration (final DMSO concentration 0.1%).

### Virus production

For second generation, self inactivating lentiviral production, LentiX293 cells were cotransfectied with pLVX transfer vector (Takara-Bio), pMD2.G (VSVg) envelope (Addgene #12259) and psPAX2 packaging (Addgene #12260) plasmids. Viral supernatant was harvested on days 2 and 3 after transfection and concentrated by ultracentrifugation for 2 h at 24,000 g (Beckman Coulter). Functional LV titers (TU/ml) were determined on Jurkat cells by FACS analysis.

### Primary human T cells culture and lentiviral transducition

Peripheral blood mononuclear cells (PBMCs) were isolated from buffy coats of healthy and anonymous donors in agreement with institutional consent and collection guidelines (Blood Transfusion Centre of Slovenia (BTC), University Hospital of Ljubljana, Permit Number: 0120-21/2020/4). PBMCs were isolated using Lymphoprep™ (STEMCELL Technologies) density gradient centrifugation and cryopreserved until further use. Isolated PMBCs were crypreserved until further use. Primary human T cells were purified from PMBCs using the Pan T cell Isolation Kit (Miltenyi Biotec). T cells were cultivated in RPMI-1640 supplemented with 10% v/v FBS and IL-2 (50 U/ml) (STEMCELL Technologies). Primary T cells were activated with anti-CD3/CD28 Dynabeads (Thermo Fisher Scientific #11132D) at a 1:1 bead-to-cell ratio. After 2 day activation human T cells were transduced with concentrated lentivirus. A cell density of 1 x 10^6^ cells/ml was maintained for expansion. The transduced T cells were adjusted for equal CAR construct expression before all downstream assays by diluting the efficiently transduced samples with untransduced T cells from the same donor. For drug wahout, the stimulated cells were wached three times in PBS and resupsended in fresh T cell media.

### Luciferase based killing assays

Luciferase-expressing target cells were plated in CoStar Black or White 96-well plates (Corning) for 24 h before adding CAR T cells, and stimulation with ligands. T cells were washed and added to target cells at the indicated target:effector (T:E) cell ratio. Co-cultures were performed in RPMI-1640 without adding IL-2. Following 24 h incubation, supernatants were harvested for ELISA, and 10 µl of prepared D-luciferin (150 ug/ml, Xenogen) was added to each well. Luminescence was measured with IVIS® Lumina Series III (Perkin Elmer) for black plates or Centro LB 963 microplate reader (Berthold Technologies) for white plates. Data were analyzed with Living Image® 4.5.2 (Perkin Elmer) or Simplicity software version 4.2 (Berthold Technologies). The viability (%) was calculated as: [1-(experimental/maximal)*100], where maximal refer to the luciferase signal from target cells alone.

### Cytokine Release Assays

Cytokines in the cell media were determined with human IFN-γ and human IL-2 ELISA kit (Uncoated ELISA Kits, Invitrogen). Briefly, primary CAR T cells T were incubated with MCF-7-fLuc or Raji-fLuc target cells with or without added ligands. After one day, the co-culture supernatant was assessed to determine IFN-γ or IL-2 levels according to the manufacturer’s instructions. Absorbance was measured at 450 nm with wavelength correction at 580 nm on a Synergy Mx automated microplate reader (BioTek) with Gen5 Microplate Data Collection and Analysis software.

### T-cell proliferation assay

To assess T cell proliferation, T cells were covalently labeled with CellTrace™ Violet-CTV (Thermo Fisher Scientific) before co-cultivation with MCF-7-fLuc target cells in RPMI 1460 media without IL-2 supplement. Five days after cultivation, the decline in CTV fluorescence was measured by FACS analysis on spectral flow cytometer Aurora with the blue, violet and red lasers (Cytek) with SpectroFlo (Cytek) software. Data was analyzed using FlowJo (TreeStar).

### Statistics and Reproducibility

Biological replicates represent parallel measurements of cells cultured in multi-well plates under the same conditions. At selected time points after cell transfection and/or stimulation, each well (biological replicate) was measured individually. . Independent experiments indicate the repetition of the whole experiment. The dose-response curves were fit to four parameter logistic (4-PL) curve using GraphPad Prism 8 (log[agonist] vs. response – variable slope; Equation: Y=Bottom + (Top-Bottom)/(1+10^((LogEC50-X)*HillSlope)). All data represents the calculated averages ± s.d. of representative data from two independent experiments. All statistical analysis was calculated with GraphPad Prism 8.

